# Functional antagonistic interactions and genomic insights into the biosynthetic potential of human gut-derived microbiota

**DOI:** 10.1101/2023.02.20.529173

**Authors:** Aehtesham Hussain, Umera Patwekar, Dattatray S Mongad, Yogesh Nimonkar, Swapnil Mundhe, Dhiraj Paul, Om Prakash, Yogesh S Shouche

## Abstract

Concerning the biological interactions within the gut microbiome, the specialized small molecules encoded by commensal microbes mediate distinct functional aspects. However, the landscape of antagonistic interactions mediated by specialized strains and their small molecules broadly remains. Here, we sought to evaluate antimicrobial interactions as a defensive contributor to gain new insights into structure-related functions or to bring the therapeutic potential of derived molecules. We elucidated the antagonistic landscape within a collection of 330 human-gut-derived commensal microbial strains cultivated from healthy human subjects. We characterized potential antagonistic strains and found a strain-specific selective inhibition contrary to common antimicrobial drugs that wipe out a broad range of species usually found in environmental microbes. Using functional and genomic approaches for accessing biologically active natural product molecules, we identified significant biosynthetic gene clusters (BGCs) encoding the important compound families in representative gut strains which contribute to antagonistic activities and are important in host defense or maintaining homeostasis in the gut. The subsets of the BGCs were represented in metagenomics sequencing data from healthy individuals. The cell culture secretome of strains revealed potential biomarkers linked to hallmark pathways. Together, these microorganisms encode biosynthetic novelty and represent a source of biologically significant natural products important in developing new treatments for infectious diseases to cut the usage of broad-spectrum antibiotics and represent a way to combat antimicrobial resistance. Consortia of such strains can be utilized as an option for precise editing of the microbiomes or fine-tuning the microbiota-modulating therapies.

## INTRODUCTION

The human microbiome encompasses a diverse network of microbial species, exhibiting composite relationships with their hosts [1, 2]. As consequences of molecular interactions that exist between the commensal microorganisms and their hosts, the microbial effects are enumerated in many ways, such as colonization resistance against the host-specific pathogens, communication, and signaling, but the precise mediators of such interactions and their complexity is largely undefined [3]. Presently more than a thousand bacterial species inhabiting the human gut support distinct functions [4]. The changes in the gut bacterial populations revealed that interactions between microbes and hosts are associated with different observations of clinical importance [5, 6]. Although contribution of gut microbiota in digestion of food, providing essential nutrients, or averting pathogens has been described, nevertheless, microbial strains regulate microbial competition in the human gut despite the gut being a frequent site for infection [7, 8]. The microbes from the human gut have a notable potential to biosynthesize and transform structurally distinct molecules [9]. Bacteria use small molecules molecules as signals to mediate their effects. These molecules directly influence host biology by targeting host cells and their receptors [10–14] or indirectly by recruiting other microbiome members collaboratively or competitively [15]. However, the functional aspects mediated by the specialized molecules encoded by human microbiome are not clearly identified.

To contextualize the microbial correlations for a specific phenotype, the identification or characterization of the microbiome-derived molecules has begun [16, 17]. Accessing the natural products by genomic information relies on biosynthetic gene clusters (BGCs), a group of genes that encode a biosynthetic pathway of a specialized molecule. DNA sequencing based metagenomics has related secondary metabolite-producing potential of the microbiome to different phenotypes, besides revealing thiopeptide BGCs as a common family of antibiotics in the human microbiome [18]. However, the broader basis of their antimicrobial capability remains to be explained at strain levels. Also, metagenomics approaches do not offer direct access to the producing microorganisms, and BGCs detected in such methods might not be active in screens, or their products may be involved in different functions beyond bacterial inhibition. To avoid such limitations, with an aim to characterize molecular mediators that regulate inter-species or microbe-host interactions, we used a culture-based approach because pure culture collections of microorganisms remain the valued reserves for examining the biosynthetic potential or accessing the natural-product molecules using bioactivity profiling or genome mining [19]. Recently, the culture-independent microbiota profiling with large-scale bacterial isolation efforts resulted in the successful cultivation of the majority of gut-derived species [20, 21]. Thus, we speculate that human gut as promising source of specialized metabolites that may serve in deciphering inter- and intra-species interactions or may facilitate the development of new antimicrobial drugs. Furthermore, the gut is frequent site for infections and a primary source of antimicrobial resistance development, thus, identifying inhibitory microbes or their molecules represents a way to minimize the spread of antibiotic resistance within the gut, either by targeting resistant infections or by reducing the colonization of multidrug-resistant pathogens [22]. The insights from the gut bacterial molecules may advance the understanding towards developing next-generation probiotics and interventions to treat or prevent infections.

With the significance of human gut microbiota strains and their derived small-molecule natural products in mediating the effects, we used functional profiling with genome-sequencing and metabolomics data to explore the biosynthetic capacity of human gut-derived strains. Analysing the biosynthetic gene clusters for accessing biologically active secondary metabolites directly from the antagonistic bacteria (strains that inhibit the growth of microorganisms) majorly avoids the discrepancies arising due to silent BGCs in various non-active strains detected by metagenomics methods. Briefly, we profiled the gut-derived facultative anaerobic collection of microbes for antimicrobial potential against pathogenic bacteria encompassing the major phyla associated with the human microbiome, including Actinobacteria, Proteobacteria, and Firmicutes, and then sequenced the genomes of representing antagonistic genera from our data. Overall, we forecast their therapeutic potential or ability to bring community changes in microbiomes for expediting the discovery process.

## RESULTS

### Inhibitory potential of gut-derived bacteria

To assess the inhibitory potential within an isolate collection of 330 morphologically distinct human gut-derived bacterial isolates, we screened the strains for inhibition potential against a panel of well-studied bacterial pathogens **(Supplementary Methods Table 1**) in antibiosis assay, and monitored that several inhibitory strains (20%; 65 strains) (**Supplementary Table S1**) with the antagonistic potential against one or multiple strains of model pathogens, the distribution of antimicrobial spectrum is given in **Fig.1a**. Of these 65 inhibitory strains, 43 strains (66%) were antagonistic against *M. luteus*, 20 strains (30%) against *S. aureus*, 19 strains (29 %) inhibited *E. coli*, 40 strains (61%) inhibited *P. aurogenesa*. 15 strains (23%) inhibited *K. pneumonia*, and 26 strains (43%) inhibited *E. faecalis*. Notably, most isolates inhibited only one to three other pathogenic strains, suggesting strain-specific or selective inhibition.

**Figure 1.**
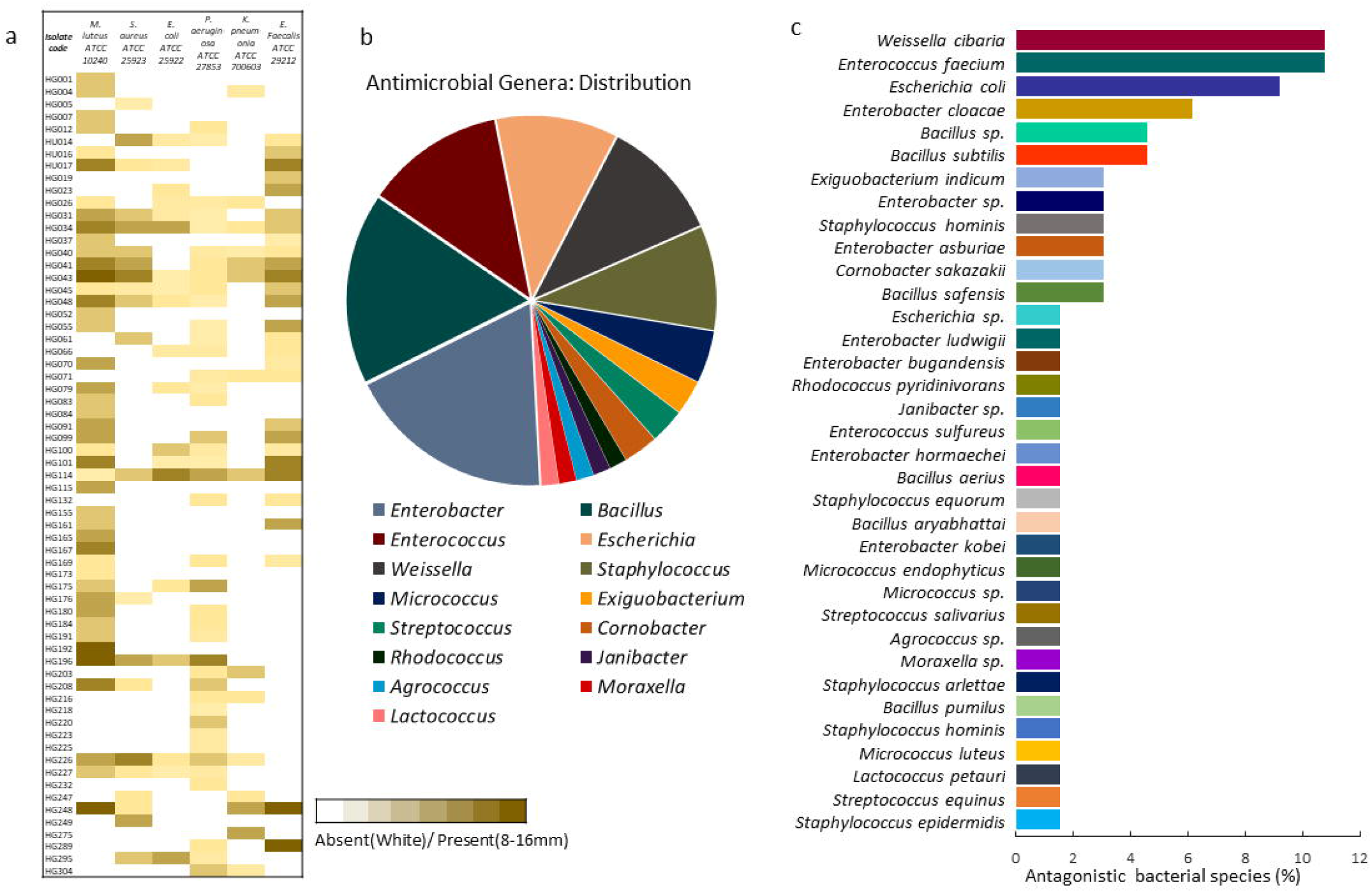
Inhibition spectrum of gut-derived bacteria. **(a)** Bioactivity spectrum of the positive hits (*n*= 65) from isolate collection of 330 gut derived microbes against human pathogens *Micrococcus luteus* (ATCC 10240), *Staphylococcus aureus* (ATCC 25923), *Escherichia coli* (ATCC 25922), *Pseudomonas aeruginosa* (ATCC 27853), *Enterococcus faecalis* (ATCC 29212) and *Klebsiella pneumonia* (ATCC 700603). The colour of the cells in the heat map represents activity zone obtained in the inhibition assay (white colour: absent; colour scale 10-16mm) **(b)** Genus level distribution of antagonistic microorganisms within the gut microbe strain library. **(c)** Species representations (%) of identified inhibitory microorganisms within collection.

Owing to the antimicrobial potential of gut isolates, we identified active strains from the isolate collection using MALDI-TOF/MS-based identification (**Supplementary Fig. S1a; Table S2**). The bacteria for which MALDI-TOF MS did not achieve reliable species-level identity, were identified using 16S rRNA gene sequencing (**Supplementary Fig. S1b; Table S3**).

Together, the MALDI-TOF MS and 16S rDNA sequencing-based identification of the microbial isolates showed that the antagonistic strains of the collection represent 35 different bacterial species belonging to 15 different genera (**Fig. 1b**, **Supplementary Table S4**) including: *Enterobacter* (18.4%; 12 strains), *Bacillus* (16.9%; 11 strains), *Enterococcus* (12.3%; 8 strains), *Weissella* (10.7%; 7 strains) *Escherichia* (10.7 %; 7 strains), *Staphylococcus* (9.2%; 6 strains), *Micrococcus* (4.6%; 3 strains), *Cornobacterium* (3%; 2 strains), *Streptococcus* (3%, 2 strain) *Agrococcus* (1.5%; 1 strain), *Rhodococcus* (1.5% 1 strain) *Exiguobacterium* (3%; 2 strains), *Janibacter* (1.5%; 1 strain), *Lactococcus* (1.5%; 1 strain), *Moraxella* (1.5%; 1 strain). However, six isolates were assigned genus levels only. In totality, all bacterial inhibitions revealed that most observed inhibitions were due to the activity of four bacterial orders Bacillales, Enterobacterales, Micrococcales, and Lactobacillales. These taxa conferred over 80% of the inhibitions of the isolate collection. In the strain collection, species-wise bacterial species *Weissella cibaria, Enterococcus faecium, Escherichia coli*, and *Enterobacter cloacae* showed high representations in inhibition assay (**Fig. 1c**). In a total of 165 observed inhibitions, a closer assessment of the top inhibiting strains revealed that *Micrococcus luteus* was susceptible to most of the gut strains. Largely, different spectrums of inhibitions were observed between the members of the same genus. Overall, the results showed that gut isolates tend to inhibit pathogens of different phylogenetic groups distinctly, showing a selective-antimicrobial pattern rather than broad-spectrum related strains. Nonetheless, the significance of such potential strains from healthy people can be their involvement in defence against pathogens through their direct inhibitory effects.

### Strain selection and growth analysis

The bacterial inhibitory tendencies might depend on the circumstances that rely on the growth or other environmental stimuli. Thus, based on the inhibition profile of the bacteria, the common genus in human gut microbiota, and the dereplication of strains, we selected eight bacteria representing different phylogenetic clades, including *Enterococcus faecium, Rhodococcus pyridinivorans, Weissella cibaria, Enterobacter sp., Micrococcus sp., Agrococcus sp., Bacillus safensis*, and *Lactococcus petauri* for the further analysis (**Fig. 2a**). To check whether these strains are robust to gut-specific media for ruling out the possibility of media influence, the growth of the individual monocultures in gut-specific synthetic media was monitored over time (0-54 hrs), and growth curves were determined. The growth pattern analysis showed that gut-specific synthetic media (**Supplementary Methods Table 2-4**) like, Gut Microbiota Medium (GMM), M17 media, and YCFA, supported the growth of bacteria (**Supplementary Fig.S2**). Therefore based on growth characteristics, selected strains were well adapted to synthetic gut-specific media besides experimental media.

**Figure 2.**
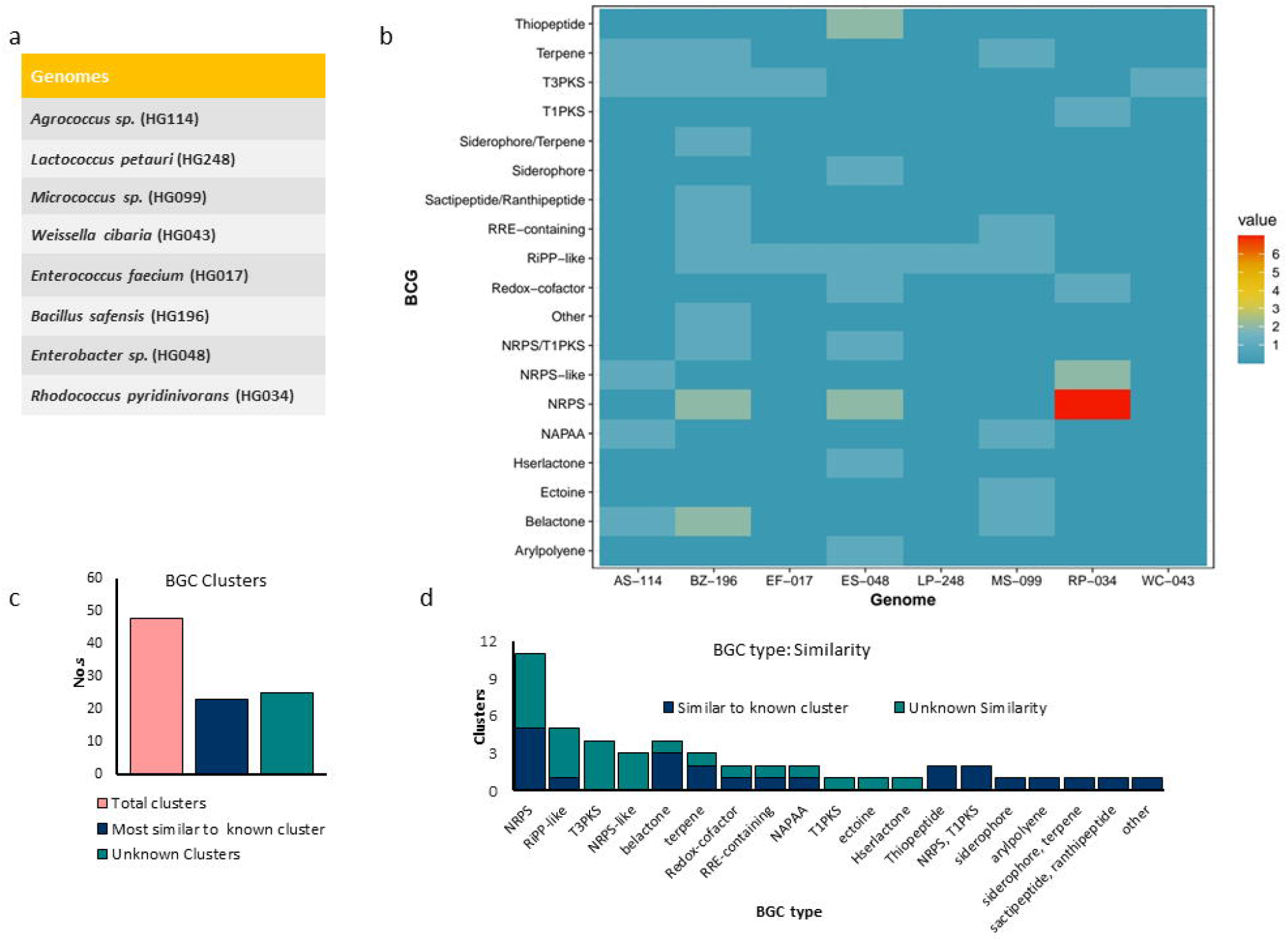
Genome annotation, identification and analysis of natural product BGCs. **(a)** List of the selected microorganisms for analysis. **(b)** Heat-map representation of number of detected BGCs of secondary metabolite families in the genomes of the bacteria *Enterococcus faecium* (HG017), *Rhodococcus pyridinivorans* (HG034), *Weissellacibaria* (HG043), *Enterobacter sp*. (HG048), *Micrococcus sp*. (HG099), *Agrococcus sp*. (HG114), *Bacillus safensis* (HG196) and *Lactococcus petauri* (HG248) are coded as EF-017, RP-034, WC-043, ES-048, MS-099, AS-114, BZ-196 and LP-248 respectively. **(c)** Bar representation of the total BGCs, similar and uncharacterized secondary metabolite BGCs in genomes. **(d)** The bar representation showing natural-product classes with characterised BGCs to that of uncharacterized BGCs that encode the biosynthetic novelty in the genomes.

### Genome sequencing, identification and analysis of natural product BGCs

The identification of vast inhibitory potencies prompted us to inspect the full biosynthetic potential of the strains. Identifying the BGCs in genome sequences has grown a potential means for natural product discovery; therefore, based on the activity profile, we sequenced genomes of eight representative bacteria. We mined the genome-sequenced strains for natural product BGCs putatively involved in the biosynthesis using Antibiotics & Secondary Metabolite Analysis Shell (antiSMASH) *[23]*. These clusters are diverse in gene content, and their biosynthetic pathways are predictable due to colocalized structure of their conserved enzymatic domains. In total, the BGCs identified in the active strains are potentially inferred to synthesize bioactive molecules belonging to distinct natural product families (**Fig. 2b, Supplementary Table S5)**.

While most of the identified molecules were encoded by distinct gut bacteria in a strain-specific manner, to identify the BGCs that likely encode the biosynthesis of uncharacterized secondary metabolite scaffolds, results from antiSMASH were compared against the database of characterized BGCs, MIBiG 2.0 [24]. The analysis revealed that 52% of BGCs are unknown BGCs, which are not connected to the MIBiG database of characterized BGCs, thus putatively encoding the biosynthetic novelty (**Fig. 2c**). Overall, BGCs in these strains belong to important natural product classes, including non-ribosomally synthesized peptides (NRPS), type I, III polyketide synthase (PKS) products, ribosomally synthesized and post-translationally modified peptides (RiPPs), terpenes, NRPS-PKS hybrids that potentially encode specialized molecules and few metabolites of uncertain function. We identified different NRPS and PKS clusters in genomes of *Agroccoccus sp*. and *Weissella cibaria*, which are not previously known to produce such classes of natural-product-like structures. Of all NRPS clusters, the majority (60%) did not cluster with any MIBiG reference entry. The T3PKS BGCs in bacteria *Enterococcus faecium, Rhodococcus pyridinivorans*, *Agrococcus sp*. and *Bacillus sp*. did not share any similarity with any MIBiG entry, which is suggestive of their biosynthetic novelty, i.e., unknown compared to the MIBiG database of characterized BGCs (**Fig. 2d**).

BGC clusters encoding RiPPs have shown a broad diversity. Only a single RiPP cluster encoding Enterocin A shows similarity to known RiPP BGCs, rest other RiPP clusters did not cluster with any of characterized RiPP BGCs, thus signifying a broad potential to identify different RiPP types, as RiPPs are the narrow-spectrum antibiotics having selective inhibition property. RiPP families: bacteriocins, lassopeptides, lanthipeptides, linaridins, microviridins and thiopeptide-linaridin hybrids might be accountable for selective inhibition against few pathogens. In addition to gene clusters for compounds involved in the inhibitory microbe-microbe interactions, BGCs potentially involved in bacterial communication, like homoserine lactones [25] were also identified.

We also detected BGCs encoding the important class of bioactive molecules like beta-lactones in many genomes and cytoprotectant molecule Ectione in the *Micrococcus sp.*. The BGCs potentially involved in metal chelation, like siderophore [26] molecules, were also detected. Thiopeptide encoding BGCs were identified in *Enterobacter sp*. As thiopeptides are modified RiPPs that possess potent antibacterial activity against Gram-positive bacteria, and currently, a semisynthetic thiopeptide LFF571 is in Phase II clinical trials for the treatment of *Clostridioides difficile* infection [27, 28]. In addition to Type-I PKS in *Rhodococcus pyrividans*, the nonribosomal peptide-polyketide synthesis hybrids (NRPS-PKS hybrids) identified in the genomes of *Bacillus* and *Enterobacter* encode important molecules like paenilamicin and saquamyacin. Also, analyses showed putative aryl polyenes in *Enterobacter sp*.

Consequently, to know the relationships among predicted BGCs, we created a BGC map to compare the degrees to which biosynthetic clusters share genes by adapting evolutionary distance between multi-domain proteins for calculating an all-by-all distance matrix [29]. In the resulting BGC distance network (**Fig. 3a**), families of secondary metabolites are widespread with respect to each other, and distinct core genetic components are more likely uniquely present in these microorganisms. In identifying the groups of related nodes, the network revealed considerable genetic variety, having groups of diverse and sparsely connected secondary metabolite biosynthetic classes, indicating that BGCs from these classes share a significant number of gene families with one another. Many of the BGCS of the particular family, which are not annotated to the characterized putative compound classes, could not subgraph with clusters of known compounds, suggesting these clusters may encode entirely different chemical scaffolds. Nonetheless, most of these inhibitory strains contain BGCs that are unique to the producing strains, and distribution specifies that the corresponding secondary metabolites could be responsible for the inhibitory role of that particular strain, as detected in the antimicrobial assay.

**Figure 3.**
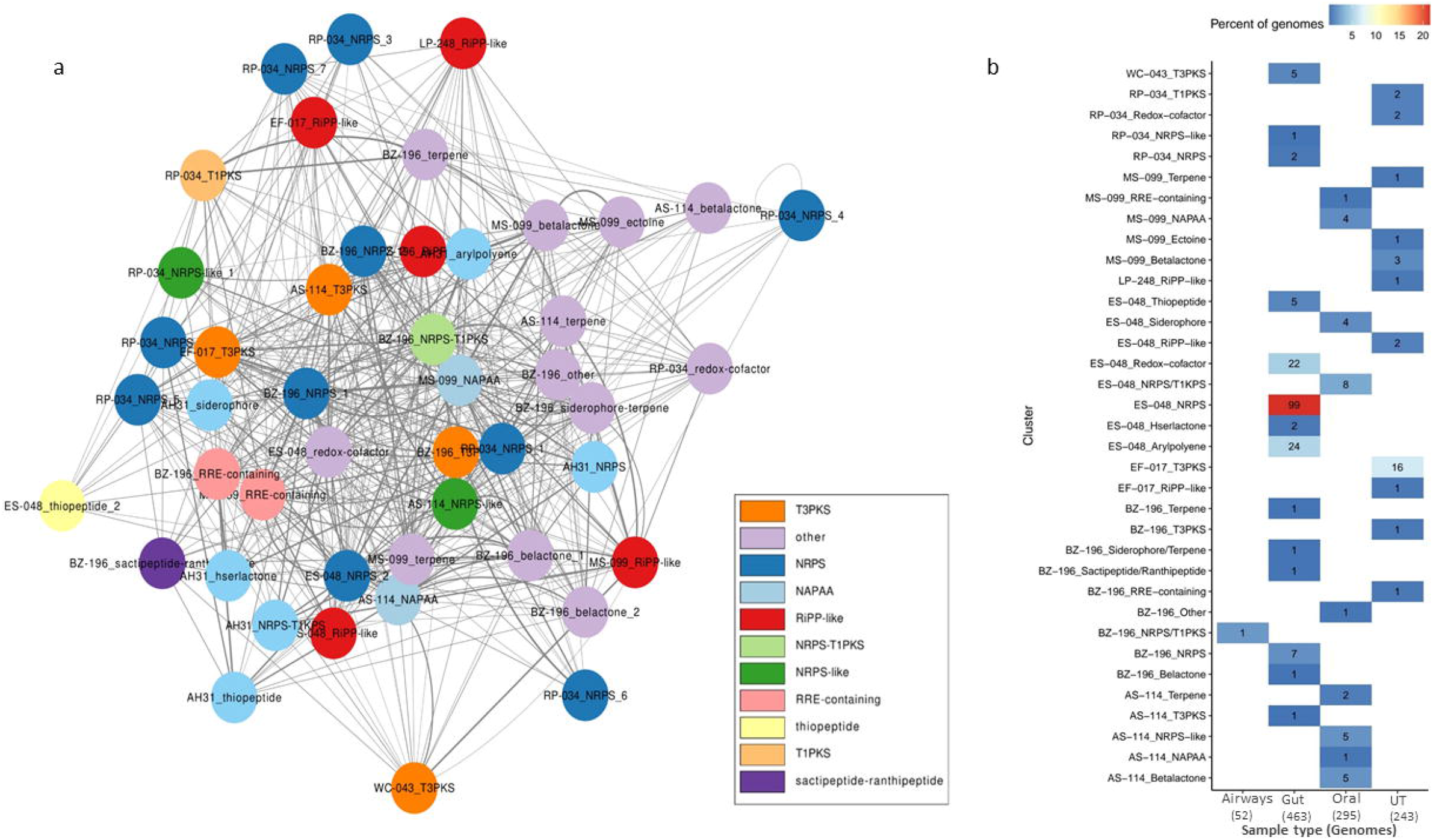
BGC analysis and BGCs in metagenomics sequencing data from healthy individuals. **a)** Gene Cluster distance Metric: Network of known and uncharacterised putative BGCs, with the BGC similarity metric threshold at 0.5. The nodes represent BGCs and clusters representing different families of natural products within genomes of *Enterococcus faecium* (HG017), *Rhodococcus pyridinivorans* (HG034), *Weissella cibaria* (HG043), *Enterobacter sp*. (HG048), *Micrococcus sp*. (HG099), *Agrococcus sp*. (HG114), *Bacillus safensis* (HG196) and *Lactococcus petauri* (HG248) are presented as EF-017, RP-034, WC-043, ES-048, MS-099, AS-114, BZ-196 and LP-248. **b)** Heat map depicting representation of BGCs amongst 1053 HMP metagenomic reconstructed genomes from four body sites (463 Gut, 295 oral, 243 UT, 52 airways). The colour of cells in the heat map represents percentage of cluster within genomes of that origin. EF-017, RP-034, WC-043, ES-048, MS-099, AS-114, BZ-196 and LP-248 represent genomes mentioned in Fig 2a.

### Representation of BGCs in the HMP genomes

Speculating these natural product encoding BGCs are involved in the microbe-host and microbe-microbe interactions, and to know whether subsets of these predicted BGCs are commonly present in healthy individuals, we tested the representation of these BGCs in the Human Microbiome Project (HMP) metagenomics sequencing data sets consisting of 1053 reconstructed genomes from different sites of the body (463 Gut, 295 oral, 243 UT, 52 airways) [30]. Most of the BGCs detected from the antagonistic gut isolates are distributed widely in healthy human-derived genomic data sets, and many of the BGCs have not been described or studied for their biosynthesis, showing the importance of natural products from such BGCs in these microbes. 172 BGC hits were found within 463 HMP genomes of gut origin, 31 among 243 urinary tract HMP genomes and 31 within 295 oral ones. In total, out of 235 hits, 172 biosynthetic gene clusters (73.19%) were present in the gut samples of healthy individuals. However, very few BGCs were found in airways genomic samples. The distribution and abundance of these BGCs (**Fig. 3b)** was predominantly in gut samples, though evident from the origin of their isolation, but confirms their overall broad representation and suggests the importance of these clusters in the microbe-host and microbe-microbe interactions among microbial inhabitants.

### Metabolomics

Subsequently, we sought to measure the metabolites secreted by these strains using liquid chromatography-mass spectrometery (LC-MS), these microorganisms were grown in broth media, and the crude cell-free supernatants were solvent extracted prior to untargeted analysis for secretory metabolites. The metabolic potential detected through LC-MS revealed another set of molecules. The analysis of the mass spectrometric data led to the detection of 603 molecules of bacterial origin uniquely present in these strains belonging to distinct compound families, including many antagonistic metabolites (**Fig 4a**; **Supplementary Table S6)**. However, 89 compounds were commonly detected in these strains, including various aminoacids, indoles, phenolics, quinones, phthalides, fatty acids, dicarboxylic acids, and other important microbial metabolites (**Supplementary Fig. S3a, Table S7**). PCA analyis of commonly detected metabolites showed a distinctive parting among the bacterial species and shown that metabolome is different from each other as samples are separated from each other (**Supplementary Fig. S3b,c).** However, overall detected metabolites cannot be precisely linked to the predicted BGC types owing to limitations associated with spectra identification libraries of known molecules. Also, most BGCs are predicted as unknown types in our genomic data sets. Nonetheless, some microbially derived compounds among the uniquely detected metabolites include important bacterial compounds involved in inhibitory and quorum-sensing phenomena like homoserine lactones and other significant molecules involved in bacterial interactions, signaling, or host physiological responses **(Supplementary Table S8**). Owing to the relevance of these strains in mediating host interactions or their administration as probiotics, we checked the relevance of detected metabolites in metabolic pathways, we found important markers in the secretome of the strains relevant to human health. While relating the metabolites to their corresponding pathways for probable functional roles revealed that many of the metabolites are involved in distinct pathways **(Fig. 4b)**, create interesting candidates for biomarkers, and suggest the importance of these bacteria or their molecules in human health.

**Figure 4.**
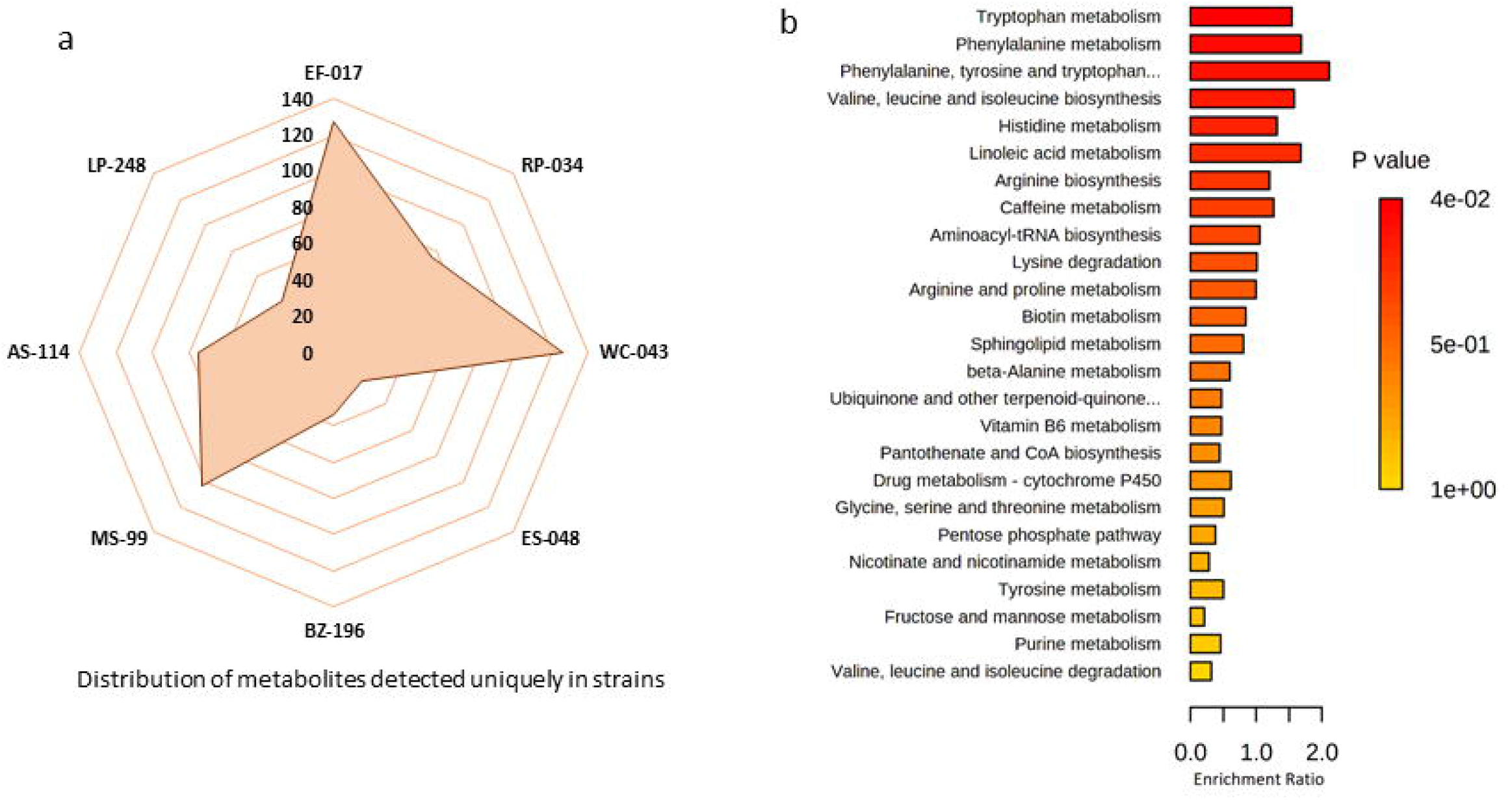
Unique metabolic features in the pathways. **(a)** A raptor representing the summary of uniquely detected metabolic features from the extracts of strains using untargeted LC-MSmetabolomics analysis. **(b)** The bar graph representing the functional role of uniquely detected metabolites from scretome of strains by LC-MS metabolomics.

## DISCUSSION

Gut microbiota interacts extensively with the host through its metabolic activities and co-metabolism of substrates. Specialized metabolites mediate a variety of cellular intraspecies and interspecies functions to perform key roles. Recently, a metagenomics-based study found a family of BGCs encoding thiopeptide as a common family of antibiotics in the human microbiome [18], However, the microbiome data are very high-dimensional, might have thousands of taxa in a sample in such analyses, and it often becomes unclear whether only one of the taxa present, or whole the community or combination of both, is responsible for phenotype [31], To overcome such fundamental barriers and assuming that the human gut as promising source of specialized metabolites of pharmaceutical importance to address the increased need for antimicrobial scaffolds, we applied the functional screening and genome mining of gut microbe collection to evaluate the potential for unprecedented antimicrobials. The functional approach categorizes the gut microbiota more precisely to reflect its suitability for human health [32]. Nevertheless, genome mining is potentially essential to increase the discovery rate and expedite the characterization of novel molecules and their biosynthetic pathways [33]. We identified different types of BGCs encoding the chemical repertoire of compound families putatively involved in bacterial defence or their niche adaptation in active strains. The BGCs detected in these bacteria belong to diverse biosynthetic classes, including NPRSs, RiPP, PKSs, terpene systems, and thiopeptides. However, most of the detected BGCs were not similar to that of characterized gene clusters, suggesting a high potential for uncharacterized molecules from such bacteria for experimental mining in drug discovery paradigms.

The BGCs containing nonribosomal peptide synthetases (NRPSs) and polyketide synthases (PKSs) are of particular interest, as they use modular enzymatic domains to synthesize molecules with complex structures, and products of their enzymes encompass several antibiotics, siderophores, antifungals and immunosuppressants [34]. Most of the BGCs from our data sets are widely distributed in healthy human samples from the different sites of origin [30], suggesting the importance of small-molecule natural products from these common BGCs.

Since, collection showed selective pathogenic inhibition, making way for potent specialized and selective metabolites. A crucial significance of such microbes would be their ability to restore microbial communities more precisely because microbial infection treatments by conventional broad-spectrum antibiotics destroy the commensal microbial communities collaterally and exert selection-pressure towards the resistant pathogenic strains which expands the number of resistance genes within the gut. However, the narrow spectrum character of strains is more appropriate for targeting specific pathogens without altering natural microbial populations, thus, can help to maintain microbial balance. This also presents a way to minimize the spread of antibiotic resistance. Since antibiotic-resistant infections pose a significant threat to human health, the gut represents the most frequent site of infection and resistance development [35]. These commensal antagonistic bacterial species can be developed into next-generation therapeutics to cut the usage of broad-spectrum antibiotics or to enhance the colonization resistance, thus can be significant in preventing resistance development which is acquired through mutation and horizontal gene transfer [36]. Also, commensal microbial species have coevolved with human hosts; thus, their safety can be considered well-established. Such molecules from antagonistic bacteria may also help to attain the defined information on the precise microbial contributions of microbiome and to fine-tune the microbiota-modulating therapies. This data provides a foundation for the relevance of such strains or their derived bioactive molecules as intervention therapies for modulating the community composition and circumventing the safety concerns associated with clinical experimental attempts like faecal transplants.

Many bacterial species from our data are studied or administrated to confer therapeutic or beneficial effects as probiotic or supplemental forms, like *E. faecium* as a probiotic in treating diarrhea [37,38], and *Lactococcus petauri* as a candidate probiotic [39]. Besides the probiotic properties, some have proven significant in clinical tails like *Weissella cibaria* [40], and *Bacillus spp*. [41]. However, our observed antimicrobial principles may serve as a proof of concept required to unlock their potential as next-generation biotherapeutics. Thus, comprehensive evaluation of these bacterial strains may bring about their employability as intervention therapies to prevent the invasion of enteric pathogens such as *Listeria, Escherichia coli* and *Salmonella*. Moreover, the secreted microbial metabolites from such strains are linked to metabolic pathways, representing the suggestive role of these bacteria or molecules in molecular mechanisms that control host-microbe interactions.

Conclusively, we uncovered the antagonistic pattern in the collection of gut microbes and explored secondary metabolite synthesis in some bacterial genomes. Although, we predict the basic chemical scaffold derived from the biosynthetic genes reported here, however a large percentage of known polyketide and nonribosomal metabolites isolated from microbial sources have antimicrobial activity [42]. Also, the observed inhibition activity of the strains has an association with specific NRPS and PKS gene clusters, suggesting that these gene clusters are involved in antibiotic production. These findings highlight the usefulness of the human gut microbes or their employability to develop new treatments for infectious diseases, open the way for complete characterization of biosynthetic genes for discovering druggable novel molecules or understanding of microbe-host interactions. Thus, we uncovered evidence for secondary metabolite synthesis in bacterial strains that have not been previously linked to such a biosynthetic repertoire and also revealed and human microbiome is a promising resource for metabolites with distinct structures.

## ACKNOWLEDGMENTS

Authors thank Director NCCS and Department of Biotechnology (DBT), Govt. of India for supporting NCMR (Grant No. BT/Coord.II/01/03/2016) at National Centre for Cell Science (NCCS), India.

## AUTHOR CONTRIBUTIONS

AH, OP, YSS designed the study. AH, UP, YN, SM performed experiments. AH and DSM analysed data. AH wrote the main manuscript. AH and DSM prepared figures, and DP, OP, YSS revised the manuscript. All authors approved the final manuscript.

## ETHICS APPROVAL AND CONSENT TO PARTICIPATE

The faecal sampling protocol was approved by the Institutional Ethical Committee and all handling procedures were performed according to SOPs.

## AVAILABILITY OF DATA AND MATERIALS

The genome sequencing data generated for the strains in this study is available in NCBI GenBank within the BioProject number: PRJNA864613.

## SUPPLEMENTARY FIGURE LEGENDS

**Supplementary Figure S1.** Identification of antagonistic bacteria.

**(a)** Dendrogram of the identified strains generated based on the MALDI-TOF/MS profiles of bacteria.

**(b)** Phylogenetic tree of the strains based on 16S rRNA gene based phylogenetic analysis, constructed by MEGA-X with neighbor-joining method using 1000 replications of the bootstrap resampling.

**Supplementary Figure S2.** Growth analysis of bacteria in gut specific media.

Growth profiles acquired by OD600 measurements of the monocultures cultures in NB and gut-specific synthetic media over 54 hrs.

**Supplementary Figure S3.** Common microbial metabolic features in bacteria.

**(a)** Heat map representing the distribution of the common microbial-derived metabolites detected in extracts of the monocultures. Extracted metabolites from monocultures of *Enterococcus faecium* (HG017), *Rhodococcus pyridinivorans* (HG034), *Weissella cibaria* (HG043), *Enterobacter sp*. (HG048), *Micrococcus sp*. (HG099), *Agrococcus sp*. (HG114), *Bacillus safensis* (HG196) and *Lactococcus petauri* (HG248) are presented as EF_17, RP_34, WC_043, ES_048, AS_114, MS_099, BZ_196, and LP_248 respectively.

**(b)** Principal component analysis plot representing common microbial-derived metabolic features from the bacterial strains.

**(c)** Dendrogram presenting the metabolic features detected commonly in secretomes of bacteria by LC-MS metabolomics.

## METHODS

### Cultivation of bacteria

The microbial strains were cultivated according to Prakash *et al*. [43]. In brief, fresh faecal samples (sampling protocol approved by the Institutional Ethical Committee) were collected from the healthy volunteers with their prior consent. The faecal samples were transported in pre-sterile containers to the laboratory at 4°C. Isolation of bacteria was carried out by serial dilution standard spread plate method using 33 different media types (for composition: **Supplementary Methods Table 6**), at 37 °C for 24-48 hrs. A total of 330 purified cultures (preserved in 20% (v/v) glycerol at −□80 °C) were reviewed for further experiments.

### Antagonistic evaluation

The antimicrobial activity against a panel of pathogenic bacteria (covering the major phyla associated with the human microbiome, including Actinobacteria, Proteobacteria, and Firmicutes species) was identified by screening 330 bacterial isolates for their capacity to inhibit the growth of pathogens. The pathogen panel included: *M. luteus* ATCC 10240, *S. aureus* ATCC 25923, *E. coli* ATCC 25922, *P. aeruginosa* ATCC 27853, *K. pneumonia* ATCC 700603, *E. faecalis* ATCC 29212. The test strains were screened in duplicate for the antibiosis potential by the spot assay and agar well diffusion assay. In spot assay, the Muller Hinton Agar (MHA) plates were inoculated by logarithmic cultures of pathogenic strains at a density of 10^5^, and test strains were inoculated on MHA plates. The plates with resulting lawn of pathogens were incubated at 37 °C for 24–48 h. In well diffusion assay, the bacterial pathogenic strains (density ~10^5^) were swabbed on MHA plates punched with wells (6 mm diameter), and 50 μL cell-free supernatant of test strains was added into the wells. The MHA plates were incubated at 37 °C for 24–48 h. Inhibition zones were measured in millimeters (mm) [44].

### MALDI-TOF/MS-based identification

MALDI-TOF/MS based identification of bacteria was carried out using alpha-cyano-4-hydroxycinnamic acid (HCCA) in standard solvent (acetonitrile: 50%, trifluoroacetic acid 2.5% and water 47.5%) as matrix. Sample preparation, ethanol/formic acid (EtOH/FA) extraction was carried as described by Kurli *et al*. [45]. Samples were analysed with Autoflex speed MALDI–TOF/TOF mass spectrometer (Bruker Daltonik GmbH, Germany) in linear positive ion extraction mode with mass cut off at m/z range: 2000–20 000Da. The MALDI Biotyper software 3.0 (Bruker Daltonik, GmbH Germany) was used to visualize mass spectra. Bacteria with biotyper score values < 2.0 were considered with species-level identity.

### 16S rRNA gene sequencing and identification of antagonistic bacteria

Genomic bacterial DNA was isolated from overnight grown cultures (LB, 37°C) by the DNA Extraction kit (DNeasy Kit, Qiagen Germany) following the manufacturer’s instructions. The amplification of 16S rDNA regions was performed with universal primers [46]. The amplified reaction products were purified using QIAquick Purification Kit (Qiagen, Germany), followed by DNA sequencing with 3730xl DNA Analyzer (Applied Biosystems). The consensus 16S rRNA gene sequences were searched against validly published bacteria available using the EzBioCloud Database [47]. The phylogenetic analysis was done via MEGA X software [48].

### Whole Genome Sequencing and Genomic assembly

High-quality DNA from the antagonistic gut strains was extracted using QIAamp DNA minikit (Qiagen, Germany) following the manufacturer’s instructions. The sequencing libraries were built from Genomic DNA using Nextera DNA Flex Library Preparation Kit with Nextera XT Index kit based on the manufacturer’s protocols (Illumina). Libraries were quality-checked and quantified using TapeStation (Agilent Technologies, USA). The quantified and denatured libraries were sequenced using paired-end 2×250-bp chemistry with 5% PhiX as an internal control on the Illumina MiSeq 2000 platform (Illumina Inc., USA). The quality-filtered reads were de-novo assembled using SPAdes (version 3.11.1). The genome assembly quality was checked by QUAST (version 5.0.2). The genome sequences of strains were submitted in NCBI GenBank (BioProject: PRJNA864613).

### Genome annotation and BGC prediction and analysis

The resulting assemblies were annotated using Rapid Annotations using Subsystem Technology (RAST) server (http://rast.nmpdr.org), and GenBank formatted files were used as input in antiSMASH 6.0 to mine the genomes for the presence of biosynthetic gene clusters of secondary metabolites [23]. BGCs from antiSMASH analyses were compared to the MIBiG database to look for their similarity with known BGCs [24].

### The BGC Map based on the gene cluster distance metric

The domain similarity index was calculated as suggested by Lin *et al*. [29]. Briefly, we derived final similarity index between two BGCs using Jaccard index: proportion of shared domains; Goodman–Kruskal γ index: similarity of the order of two distinct domains between two BGCs; and Domain duplication similarity index: amount of duplication of a shared domain as described.

### Representation of BGCs in the HMP genomes

To know the representation of BGCs in the HMP metagenomics-sequencing data from healthy humans, BGCs were inquired against 1053 reconstructed genomes from different sites of the body, including 463(Gut), 295 (oral), 243 (UT), and (52) airways. These HMP genomes were downloaded from: https://www.hmpdacc.org/hmp/catalog/ [30].

### Growth analysis in gut-specific media

To determine the growth of bacteria in gut-specific synthetic media, growth was monitored over time in 4 different media, including NB and three gut-specific media (YCFA, GMM, and M17) (for composition: **Supplementary Methods Table 2-4**). Bacterial inocula were prepared from freshly grown bacterial monocultures and diluted in respective media to 0.01 OD_600_nn. The growth was measured in 96 well round-bottom plates (Himedia) in triplicates at time intervals of 0 hrs, 8 hrs, 24hrs, 32hrs, 48 hrs, and 54 hrs using a SpectraMax Plus 384 Microplate Reader (to record absorption wavelength at 600□nm).

### Bacterial fermentation and LCMS sample preparation

Individual colony from each monoculture was inoculated in nutrient broth and incubated at 37°C with shaking at 110 rpm to generate seed cultures. The seed cultures were used to inoculate broth cultures in 1000-ml Erlenmeyer Flasks. After 2-4 days, supernatants from the bacterial cultures were extracted with 2x ethyl acetate. The ethyl acetate layers were concentrated in vacuo, and re-dissolved methanol. The concentrated extracts were filtered with 0.2 □μm amicon filter (Millipore), quantified and examined by LC-MS analysis.

### Mass spectrometry and untargeted metabolomic analysis

Untargeted metabolomics profiling was performed using Q-Exactive-Orbitrap mass spectrometer (Thermo Fisher) coupled online with Accela™ 1250 ultra-high performance liquid chromatography (UHPLC) system (Thermo Fisher) via heated electrospray ionization source (HESI) operated in positive ion mode. Raw profile mode data files were converted to mzXML open format and subjected to baseline correction, chromatogram alignment, peak picking and feature annotation using packages like ProbMetab, xcms, CAMERA in R environment (version 3.3.3). Putative identification of metabolites was carried by comparing annotated features to KEGG database (http://www.kegg.com). Multivariate statistical analysis was performed by MetaboAnalyst 5.0 [49]. For pathway analysis, correlations between microbes and metabolites were obtained using Pearson correlation coefficient.

