## Supplementary figures and images for "Functional antagonistic interactions and genomic insights into the biosynthetic potential of human gut-derived microbiota"

### Supplementary Figure S1

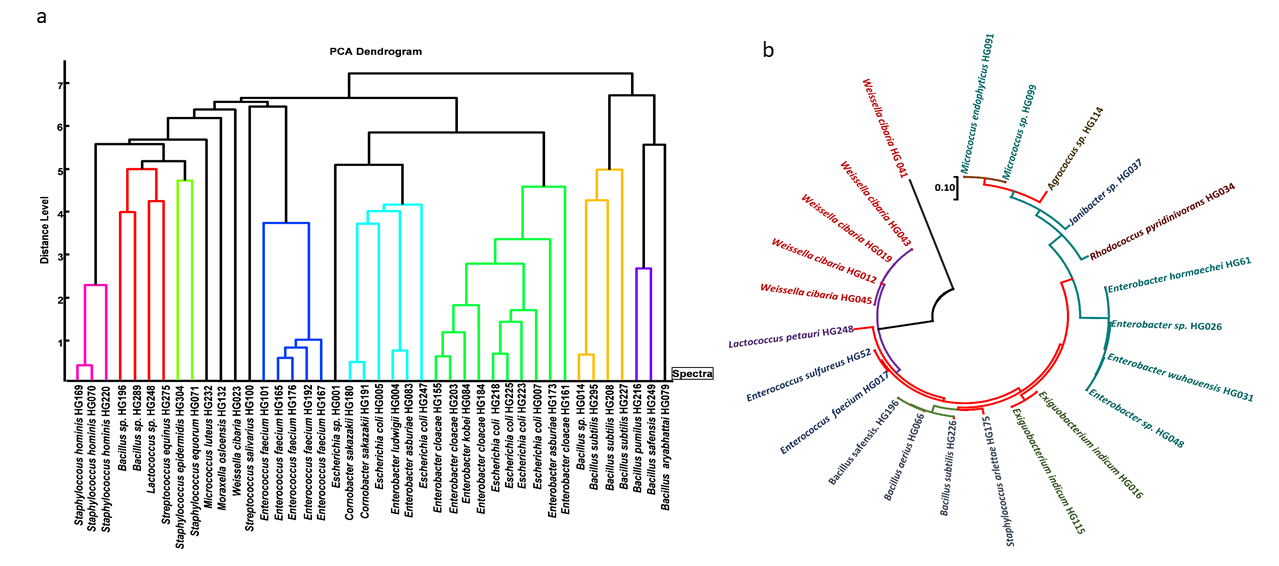

### Supplementary Figure S2

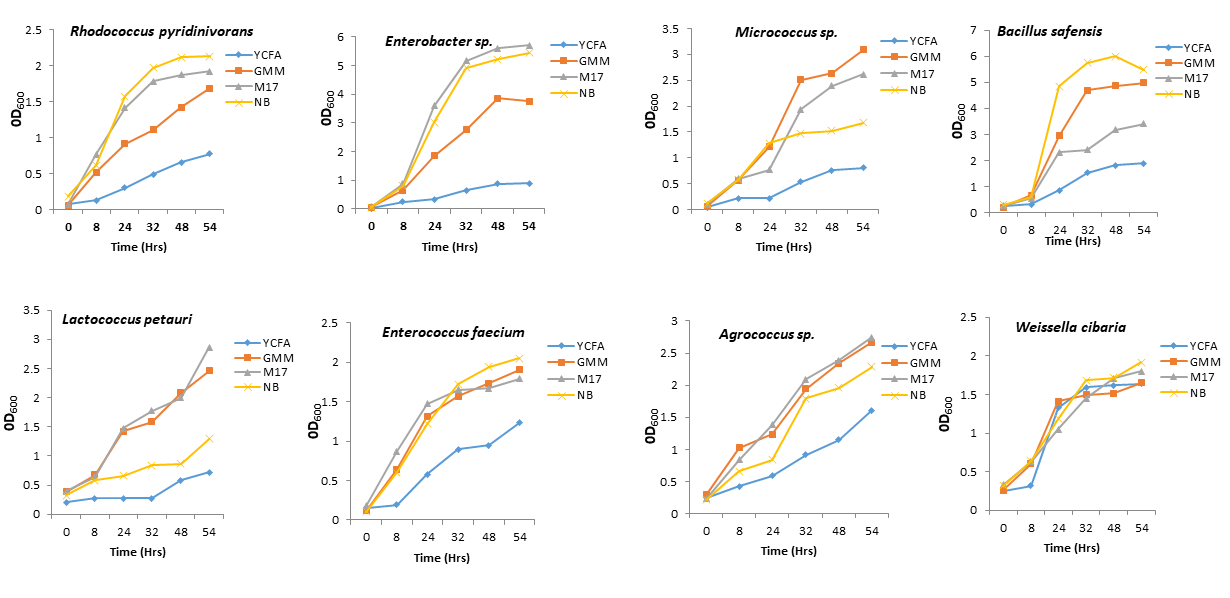

### Supplementary Figure S3

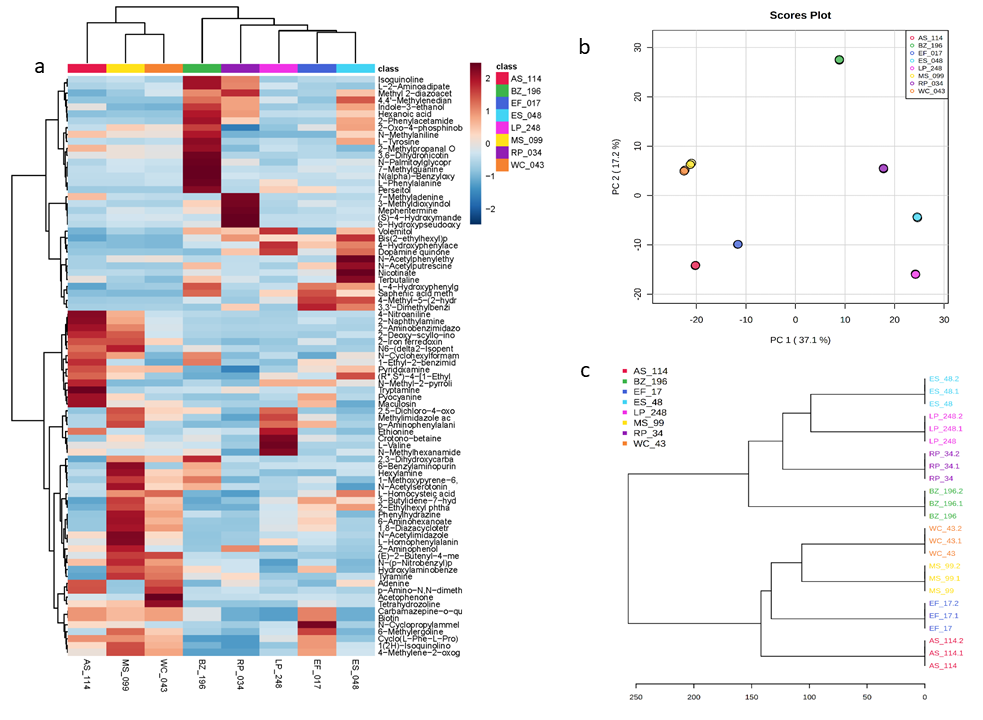
