## Supplementary Methods for "Functional antagonistic interactions and genomic insights into the biosynthetic potential of human gut-derived microbiota"

**Table 1:**

**Pathogenic strains:**

| **S. No.** | **Pathogen** |
| --- | --- |
| 1 | *Staphylococcus aureus* (ATCC 25923) |
| 2 | *Micrococcus luteus* (ATCC 10240) |
| 3 | *Escherichia coli* (ATCC 25922) |
| 4 | *Pseudomonas aeruginosa* (ATCC 27853) |
| 5 | *E. faecalis* (ATCC 29212) |
| 6 | *Klebsiella pneumonia* (ATCC 700603) |

**Table (2-4): Synthetic media for used for growth analysis of microbes**

**Table 2. Gut Microbiota Medium – GMM**

| **Components** | **Amount/ L (Concentration)** | **Comments** |
| --- | --- | --- |
| Tryptone peptone | 2 g |  |
| Yeast extract | 1 g |  |
| D-glucose | 0.4g |  |
| Maltose | 1g |  |
| Fructose | 1g |  |
| Cellobiose | 1g |  |
| L – cysteine | 0.5 g |  |
| Meat Extract | 5 g |  |
| KH_2_PO_4_ | 100 mL (100mM) | 1M stock solution pH 7.2 |
| MgSO_4_-7H20 | 0.002 g |  |
| NaHCO_3_ | 0.4 g |  |
| NaCl | 0.08 g |  |
| CaCl_2_ | 1 mL | 0. 8g/100mL stock |
| Vitamin K (menadione) | 1 mL (5.8 mM) | 1 mg/mL stock |
| FeSO_4_ | 1 ml (1.44 mM) | 0.4 mg FeSO4/mL stock |
| Histidine Hematin Solution | 1 mL (0.1%) | 1.2 mg hematin/mL in 0.2M histidine |
| Tween 80 | 2 mL (0.05%) | 25% stock solution |
| ATCC Vitamin Mix | 10 mL (1%) |  |
| ATCC Trace Mineral Mix | 10 mL (1%) |  |
| Resazurin | 4 mL (4Mm) | 0.25 mg/mL stock solution |
| isobutyrate, | 4 mM |  |
| isovalerate | 1mM |  |
| propionate | 8Mm |  |
| Acetic acid | 30mM |  |

**Reference**: Goodman AL, Kallstrom G, Faith JJ, Reyes A, Moore A, Dantas G, Gordon JI. Extensive personal human gut microbiota culture collections characterized and manipulated in gnotobiotic mice. Proceedings of the National Academy of Sciences. 2011, 12; 108:62527.

**Table 3.**

**YCFA Medium**

Reference - Das P, Ji B, Kovatcheva-Datchary P, Bäckhed F, Nielsen J. In vitro co-cultures of human gut bacterial species as predicted from co-occurrence network analysis. PLoS One. 2018; 13:e0195161. [https://doi.org/10.1371/journal. pone.0195161](https://doi.org/10.1371/journal.%20pone.0195161)

| **Components** | **g/100 ml** |
| --- | --- |
| Yeast extract | 0.25 g |
| Casitone | 1 g |
| K2HPO4 | 0.045 g |
| KH2PO4 | 0.045 g |
| NaCl | 0.09 g |
| (NH4)2SO4 | 0.09 g |
| MgSO4 7H2O | 0.009 g |
| CaCl2 | 0.009 g |
| Acetate | 33 mM |
| Propionate | 9 Mm |
| isobutyrate, isovalerate, and valerate | 1 mM each |

The ingredients were boiled to dissolve completely. After cooling, solution was supplemented with

| L-cysteine | 0.1 g |
| --- | --- |
| NaHCO3 | 0.4 g |
| Hemin | 1 mg |

After autoclaving at 120˚C for 15 min, filter sterilized solutions of fallowing vitamins were added:

| Biotin | 1 μg |
| --- | --- |
| Cobalamin | 1 μg |
| p-aminobenzoic acid | 3 μg |
| folic acid | 5 μg |
| pyridoxamine | 15 μg |
| Thiamine | 5 μg |
| Riboflavin | 5 μg |
| final pH | 7.2 ± 0.1 |

**Table 4.**

**M17 Medium**

| **Components** | **g/L** |
| --- | --- |
| Peptone type I | 2.5 g |
| Casein enzymic hydrolysate | 2.5 g |
| soya peptone (papainic) | 5 g |
| Yeast extract | 2.5 g |
| Beef extract | 5 g |
| Lactose | 2.5 g |
| Ascorbic acid | 0.5 g |
| Disodium β-glycerophosphate pentahydrate | 19 g |
| Magnesium sulphate | 0.25 g |
| Final pH ( at 25°C) | 7.1±0.1 |

**Table 5.**

**Nutrient Broth (HiMedia)**

| **Components** | **Gms/Litre** |
| --- | --- |
| Peptone | 10 g |
| Beef extract | 10 g |
| Sodium chloride | 5 g |
| pH | 7.3±0.1 |

**Table 6. Media used for isolation of microbes.**

| **Medium** | **Components- Gms/Litre** |
| --- | --- |
| HK1 | Baker’s yeast, soluble form- 5.00  Calcium chloride. 2H_2_O- 1.36  Agar- 15.00  Final pH (at 25°C) 7.2 ± 0.2 |
| HK2 | Yeast Nitrogen Base (M139) - 5.58  Casein acid hydrolysate- 0.0083  Dipotassium hydrogen phosphate- 1.66  Sorbitol- 10.00  Agar - 15. |
| HK3 | Glucose- 4.00  Yeast extract- 4.00  Malt extract- 10.00  Agar- 20.00  Final pH (at 25°C) 7.2 ± 0.2 |
| HK4 | Dipotassium hydrogen phosphate- 5.12  Potassium dihydrogen phosphate- 1.50  Ammonium chloride - 0.30  Calcium chloride -0.01  Magnesium sulphate. 7H_2_O- 0.20  Potassium nitrate- 2.00  Sodium benzoate- 0.92  Trace element solution SL-6- 2.00ml  Agar- 15.0  Final pH (at 25°C) 8.2 |
| HK5 | Pancreatic digest of casein- 17.00  Enzymatic digest of soyabean meal- 3.00  Sodium chloride- 13.50  Dextrose- 2.50  Dipotassium phosphate - 2.50  ACES buffer- 3.60  Agar- 15.00  Final pH (at 25°C) 6.8 ± 0.1 |
| HK6 | Peptic digest of animal tissue- 20.00  Lactose- 10.00  Bile salt sodium taurocholate- 5.00  Sodium chloride- 5.00  Neutral red- 0.07  Agar- 20.00  Final pH (at 25°C) 7.4 ± 0.2 |
| HK7 | Soil extract- 100 ml  Malt extract- 5.00  Agar - 20.00 |
| HK8 | L-Asparagine- 0.10  Dipotassium hydrogen phosphate- 0.50  Ferrous sulphate- 0.001  Magnesium sulphate- 0.10  Sodium caseinate- 2.00  Sodium propionate- 4.00  Agar- 15.00  Final pH (at 25°C) 8.1 ± 0.2 |
| HK9 | Sodium chloride- 156.00  Magnesium chloride, 6H_2_O- 13.00  Magnesium sulphate, 7H_2_O- 20.00  Calcium chloride, 2H_2_O - 1.00  Potassium chlorid - 4.00  Sodium carbonate- 0.20  Sodium bromid - 0.20  Yeast extract- 5.00  Glucose- 1.00  Agar - 15.00  Final pH (at 25°C) 7.0 ± 0.2 |
| HK10 | Ammonium sulphate- 1.00  Dipotassium hydrogen phosphate - 1.00  Magnesium sulphate. 7H_2_O- 0.20  Calcium chloride- 0.01  Ferric chloride- 0.02  Cellulose powder-10.00  Agar- 10.00  Cellobiose - 0.342 |
| HK1 | Yeast extract- 2.00  Tryptone- 1.00  Sodium acetate- 1.00  Soil extract- 50 ml  Agar 15.00  Final pH (at 25°C) 7.4 ± 0.2 |
| HK1 | Peptone- 5.00  Meat extract- 3.00  Agar- 15.00  Final pH (at 25°C) 7.1 ± 0.1 |
| HK13 | Glucose- 4.00  Yeast extract- 4.00  Malt extract- 10.00  Calcium carbonate- 2.00  Agar- 12.00  Final pH (at 25°C) 7.2 ± 0.2 |
| HK14 | L-Asparagine- 1.00  Dipotassium hydrogen phosphate- 1.00  Ferrous sulphate. 7H_2_O- 0.001  Zinc sulphate.7H_2_O- 0.001  Manganese chloride. 4H_2_O- 0.001  Agar 15.00  Final pH (at 25°C) 7.2 ± 0.2 |
| HK15 | Peptone- 10.00  Beef extract- 10.00  Yeast extract- 5.00  Glucose- 20.00  Disodium hydrogen phosphate- 2.00  Sodium acetate- 5.00  Tri-ammonium citrate- 2.00  Magnesium sulphate- 0.20  Manganese sulphate- 0.20  Tween- 80 1 ml  Agar- 15.00  Final pH (at 25°C) 6.4 ± 0.2 |
| HK1 | Soya bean- 20.00  Mannitol- 20.00  Agar- 20.00  Final pH (at 25°C) 7.2 ± 0.2 |
| HK17 | Yeast extract- 5.00  Peptone- 3.00  Mannitol- 25.00  Agar- 12.00 |
| HK18 | Peptone- 15.00  Chromogenic mixture- 2.45  Agar- 15.00  Final pH (at 25°C) 6.8 ± 0.2 |
| HK17 | Dipotassium hydrogen phosphate- 1.77  Potassium dihydrogen phosphate- 0.680  Sodium chloride- 0.140  Calcium chloride- 0.132  Magnesium sulphate- 7H_2_O 0.200  Mannitol- 10.00  Yeast extract- 0.080  Ferrous sulphate. 7H_2_O- 0.0025  Boric acid- 0.0029  Cobaltous sulphate, 7H_2_O- 0.0012  Copper sulphate, 5H_2_O- 0.0001  Manganese chloride, 4H_2_O 0.00009  Sodium molybdate, 2H_2_O 0.0025  Zinc sulphate, 7H_2_O 0.0012  Agar 15.00  Final pH (at 25°C) 7.0 ± 0.2 |
| HK20 | Malt extract- 40.00  Ox-bile (dessicated)- 20.00  Tween 40- 10.00  Agar 15.00 |
| HK21 | Casein enzymic hydrolysate- 5.00  Yeast extract- 1.00  Agar- 15.00  Final pH (at 25°C) 7.0 ± 0.2 |
| HK22 | Tryptone- 10.00  Proteose peptone- 10.00  Dipotassium hydrogen phosphate- 1.50  Magnesium sulphate- 1.50  Agar- 15.00  Final pH (at 25°C) 7.2 ± 0.2 |
| HK23 | Peptone from casein- 15.00  Peptone from soymeal- 5.00  Sodium chloride- 5.00  Agar- 15.00  Final pH (at 25°C) 7.2 ± 0.2 |
| HK24 | Peptic digest of animal tissue- 5.00  Beef extract- 1.50  Yeast extract- 1.50  Sodium chloride-5.0  Agar 15.00  Final pH (at 25°C) 9.7 ± 0.2 |
| HK25 | Ammonium sulphate- 1.00  Dipotassium hydrogen phosphate- 5.00  Fumaric acid- 3.00  Sodium formate 3.00  Yeast extract 1.00  Magnesium chloride. 6H_2_O- 0.20  Ferrous sulphate.7H_2_O- 0.02  Sodium thioglycollate- 0.50  Resazurin- 0.001  Agar 15.00  Final pH (at 25°C) 7.1 ± 0.1 |
| HK26 | Tryptone- 10.00  Sodium chloride- 24.00  Magnesium chloride. 6H_2_O- 11.00  Sodium sulphate- 4.00  Calcium chloride.6H_2_O- 2.00  Potassium chloride- 0.70  Potassium bromide- 0.10  Boric acid- 0.03  Sodium thiosulphate. 9H_2_O- 0.005  Strontium chloride.6H_2_O- 0.04  Sodium fluoride- 0.003  Ammonium nitrate- 0.002  Ferric phosphate, 4H_2_O 0.001  Agar-15.00  Final pH (at 25°C) 7.8 ± 0.2 |
| HK27 | Sodium aspartate- 10.00  Yeast extract- 0.20  Magnesium sulphate.7H_2_O- 1.00  Calcium chloride.2H_2_O- 0.028  Dipotassium hydrogen phosphate- 0.75  Sodium dihydrogen phosphate- 0.25  Cysteine hydrochloride- 0.25  Resazurin 0.001  Trace element solution 1.00ml  Agar 15.00  Final pH (at 25°C) 7.0 ± 0.2 |
| HK28 | Potassium phosphate-15.00  Ammonium sulphate- 1.00  Magnesium sulphate- 0.20  Calcium chloride- 0.01  Infusion broth- 25.00  Dextrose- 5.00  L-Cysteine hydrochloride- 1.00  Pancreatic digest of casein- 4.00  Yeast extract- 5.00  Soluble starch- 1.00  Agar- 15.00  Final pH (at 25°C)- 6.9 ± 0.2 |
| HK29 | Peptone- 10.00  Sodium chloride- 5.00  Calcium chloride.2H_2_O- 0.10  Agar- 15.00 |
| HK30 | Sucrose- 1.00  Casein enzymic hydrolysate- 0.75  Yeast extract- 0.25  Pancreatic digest of casein- 0.154  Papaic digest of soyabean meal- 0.027  Sodium chloride- 0.045  Dipotassium hydrogen phosphate- 0.022  Agar 15.00  Final pH (at 25°C) 7.1 ± 0.2 |
| Luria Agar | Tryptone-10.000  Yeast extract- 5.00  Sodium chloride- 5.00  Agar- 15.00  Final pH ( at 25°C) 7.0±0.2 |
| R2A | Casein Enzymic hydrolysate- 0.250  Peptic digest of animal tissue- 0.250  Casein Acid hydrolysate-0.500  Yeast extract- 0.500  Glucose- 0.500  Starch soluble- 0.500  Dipotassium phosphate- 0.030  Magnesium sulphate. heptahydrate- 0.500 Sodium pyruvate- 0.030  Agar- 15.000  Final pH ( at 25°C) 7.2±0.2 |
| Tryptic Soya Agar | Tryptone- 17.000  Soya peptone- 3.000  Sodium chloride- 5.000  Dextrose- 2.500  Dipotassium hydrogen phosphate- 2.500  Agar- 15.000  Final pH ( at 25°C) 7.3±0.2 |
